# Study on the Inflammatory Response of PMMA/Polystyrene/Silica Nanocomposite Membranes for Drug Delivery and Dental Applications

**DOI:** 10.1101/499160

**Authors:** Shanmuga Sundar Saravanabhavan, Kannan Natarajan, Sundaravadivel Elumalai, Sarang Zsolt, Mukunthan K Selvam, Manokaran Janakiraman, Narendranath Jonna, Kamalakannan Vasantha Palaniappan, N Kavitha, Prabhu Venkatesan, Balasubramanian Natesan

## Abstract

**Background:** The application of polymeric materials in medical industry has grown drastically in the last two decades due to their various advantages compared to existing materials. The present research work emphases on the sol-gel technique to formulate the polymethyl methyl acrylate/polystyrene/silica composite membrane.

**Methods:** The characteristic of the composite was investigated through modern state art of instrumentation.

**Results:** The functional groups attached to the polymer was absorbed by FTIR. The FTIR spectrum confirm that the blend was mixed thoroughly and the formation of unite intimately between the polymers. The membranes were observed by SEM for its surface homogeneity which depends upon the composition of the two blending polymers. The captured SEM images showed the formation of microcracks on the surface, which was evidently controlled by varying the constituent polymer ratios. The prepared blend membranes with 2:1 ratio of PMMA/PS/Si displayed higher water uptake compared to other blended membranes. The composite membranes had good hydroxyl apatite growth in SBF solution. Furthermore, the *in vitro* cytotoxicity studies carried out by MTT method, using RAW macrophage cells showed that all the samples exhibited excellent cell viability.

**Conclusion:** The inflammatory response of composite with equal concentration of PMMA-PS were performed and observed no inflammation in comparison with control and other tested concentrations.

## Introduction

Immense research in biomaterials used for hard and soft body tissues replacement and orthopedic applications were constantly increasing during the past few decades[1–5]. Biomaterials, used for this kind of replacement should be inert, bioactive and biocompatible. Based on the type of implant needed, the type of material like metals, alloys, ceramics and polymeric materials can be selected as a suitable biomaterial [6]. Among various kinds of biomaterials, polymeric biomaterials have gained more importance in recent days due to its vast advantages. Moreover, polymeric biomaterials are being used as a replacement of metallic materials (amalgam) due to their added advantage like light weight and tailor made properties[6]. There are several natural polymers like chitosan, cellulose having bioactivity and biocompatibility are used as biomaterials for various wide range of biomedical applications[7, 8]. However, the applications of these natural polymers are limited in terms of its stability and strength. The synthetic polymer is an alternative to natural polymers which may enhance the stability, strength and biocompatibility. These tailor-made properties of synthetic polymers improve its medical and biomedical applications. There is list of synthetic polymers such as polyamides (PAm), polyethylene (PE), polyether ether ketone (PEEK), poly methyl methacrylate (PMMA), polysulfone (PSu), polytetrafluoroethylene (PTFE), polyurethane (PU), and ultra-high molecular weight polyethylene (UHMWPE) used as biomaterials with inorganic nano bio-materials to induce bioactivity. The bioactivity of these polymers were induced by functionalizing the polymer or by blending with other polymer having bioactivity[9].

The PMMA is the most successful and investigated material in medical application such as implant in orthopedic applications due to its good bioactive and biocompatibility nature when used as bone cements in hard tissue replacements. Despite the many drawbacks like brittleness, shrinkage and high polymerization exotherm it used as bone cement for orthopedic applications is still serves a greater advantage. Similarly, polystyrene is one of the highly researched topic in biomedical applications known for its inert nature and has its applications in consumer, food packing industries has shown a good bioactivity and enhanced cell adhesion when modified with silica [10, 11]. It is an important criterion to use porous bioactive polymer matrix since it is more advantageous considering the ability of hydroxyapatite to grow within the pores, this phenomenon affixes the formation of interlock with the adjacent normal bone thereby lifts the fixation of polymer prosthesis. The application of these polymer matrix as biomaterials greatly influences its biological and mechanical properties in internal bone implants due to the tailor-made nature [12]. Regardless of advantages of PMMA possess a major drawback that is causing inflammation after the removal of the prosthesis from the implant site [13–14] will be addressed in the present study.

The sol-gel technique has been successfully used for preparation of PMMA/Polystyrene/Silica nanocomposite membrane. The sol-gel technique is selected due to various advantages includes, commendable chemical homogeneity, controlled size and morphology, operates at low temperature [15–17]. The sol-gel composed membranes are highly embodied matrices, which are used in the fields of biomedical sensors, laser materials and for sustained drug delivery applications. The present study propose about the bled formation between PS and PMMA based on the dipole interaction between aromatic groups of PS with that of the carbonyl group in PMMA. The schematic representation of the same is given in the Fig. 1.

**Figure 1.**
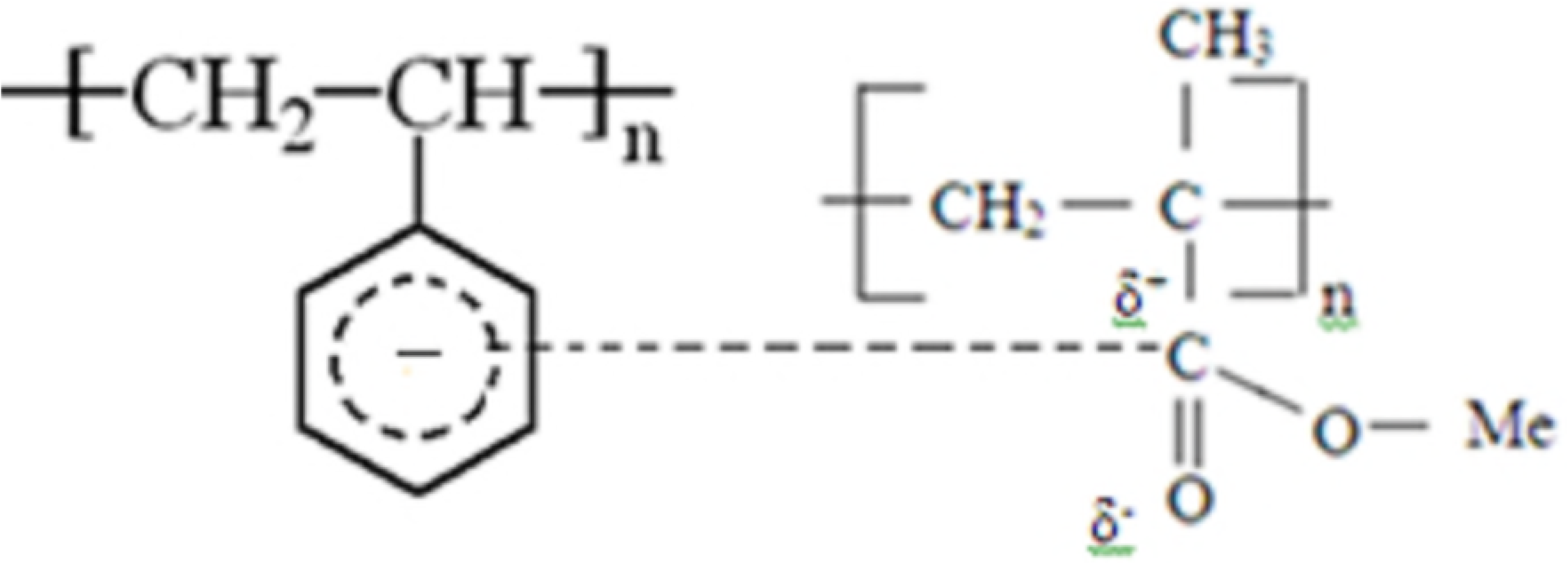
The mechanism of interaction that occurs between PS and PMMA.

Based on the literature survey and to the best of our knowledge we are the first to report about the combination of PMMA/PS/Si composite material for medical application. The silica was selected as filler material due to its biocompatibility and ease availability in nature. There were quite an enormous amount of research that have been conducted with silica based polymeric materials for biomedical applications such as implants and artificial skin [18, 19]. Apart from its biocompatibility nature, its incorporation also increase the strength of polymeric materials and influence the creation of apatite formation thereby enhancing the bioactivity of composite by interacting with negatively charged silanol and SBF solution. In the present study, PMMA/PS/Si composite membranes were prepared in different weight ratios of PMMA and PS (1:1, 1:2, 2:1) with constant weight of silica. The prepared composite membranes were tested for its bioactivity by immersing in SBF and cytotoxicity studies by MTT method.

## Materials and Methods

The materials that were used for the study were procured commercially from different sources. PMMA (Mol. Wt 35,000 Da) was procured from Asian acrylates, Mumbai, Polystyrene from Sigma Aldrich. Silica and Tetraethyl orthosilicate (TEOS, 99.5% pure) were purchased from Sigma Aldrich. THF, Hydrochloric acid was purchased from Merck. Different polymer system employed and their corresponding code, weight ratio is shown in Table 1, in all the case silica concentration was kept constant.

### Preparation of Composite membranes

The polymer silica composite membranes were prepared by weighing different amounts of PMMA and Polystyrene in separate beakers followed by dissolving in THF solvent. The polymers solutions were mixed together with constant stirring to form PMMA/PS blends. The silica particles as mentioned in Table 1 were added to the homogenous polymer solution and subsequently stirred, ultra-sonicated for 2 hours to get uniform dispersion of silica particles. In order to enhance the gelation process 0.25 ml of 35% HCl, 5 ml ethanol and 1ml of tetra ethoxy silane (TEOS) were added to the polymer solution. The solutions were left for three days stirring to complete the gelation process. The gel obtained was then casted onto clean-dried petri dishes and left undisturbed for three days at room temperature. After complete removal of solvent, the membranes were removed from the petri dish for the further analysis.

### Fourier Transform Infra-Red Spectroscopy (FTIR)

The presence and interaction of different functional groups in polymers were studied using FTIR. The composite membranes of PMMA/PS prepared were characterized using Perkin Elmer Spectrum RXI IR spectrophotometer.

### Scanning Electron Microscope (SEM)

The surface morphology and dispersion of silica fillers in the prepared composite membranes were studied using HITACHI S-3400N Scanning Electron Microscope (SEM). Prior to the analysis, the samples were dried and their surface was gold sputtered.

### Water absorption

Water absorption property of the polymer composite membranes was carried out as per our previous studies[20]. The study was carried out by immersing pre-weighed dried membranes in deionized water on a glass beaker for 24 hours. After 24 hours, wet membranes were retrieved from the beaker and excess water was blotted gently using a tissue paper. The wet samples were again weighed, and the water absorption percentage was calculated using the formula.

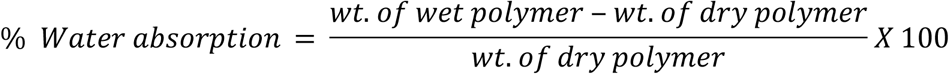

### Bioactivity Study

The bioactivity of the membranes prepared was studied by immersing the membranes in simulated body fluid (SBF) for 15 days (with and without silica fillers) and analyzing their surface for the formation of mineral (hydroxycarbonate apatite) layer using SEM (Quanta 200 FEG scanning electron microscope). SBF also known as Kokubo’s Solution was prepared according to the specification given by Kokubo et al [21]. The chemicals required to prepare SBF were dissolved in deionized water in the specified quantity while the pH was maintained at 7.25.

### Drug Release Kinetics

The effective drug release kinetics of composite membranes prepared were tested using 5-Fluorouracil as model drug. The solution was prepared by dissolving 12 mg 5-FU in 1000ml of deionized water. The composite membranes prepared were weighed and cut into 1cm^2^ followed by immersion in drug solution for 24 h. The increase in weight of sample immersed after 24 h was measured and compared with initial weight to determine amount of drug loaded onto the membrane. The amount of drug released from composite was evaluated using a UV-Visible spectrophotometer (T90+UV/Vis Spectrometer, PG Instruments) at a wavelength of 293 nm. The drug release kinetics was determined from the drug absorbance values after immersion in PBS solution and comparing it with the standard values obtained in the initial composites [22].

### Biocompatibility test

The composites prepared must be biocompatible such that it can be applied *in vivo.* Hence, in the present study we used MTT assay to quantify the biocompatibility of prepared composites. The cell viability was determined based on our previous studies [23]. The biocompatibility analysis was performed using RAW macrophage cell line 264.7 that were grown on Dulbecco’s Modified Eagle’s Medium in a 96 well plates. The cells were allowed to settle after which the samples were added and the viability ratio was analyzed based on absorbance values at 545nm. The obtained values were plotted against the standard control group and viability ratio were estimated.

### *In vitro* inflammatory studies

We have evaluated the inflammatory response of PMMA-PS composited using RAW 264.7 macrophage cells in vitro. The cells were tested with composite prepared and inflammatory responses for cytokines TNF-α, IL-6 and IL-1were checked using qPCR. Briefly, the cells were treated with the samples and allowed for 6h incubation. The first inflammatory response was analyzed by extracting the RNA and reverse transcribing the same to form cDNA. Followed by the analyzing of expression of inflammatory markers using qPCR.

### Statistical Analysis

All the experiments were repeated for five times and average from them was used for the study.

## Results and Discussion

### FTIR

The FT-IR Spectra showed the shifting of functional groups towards the lower frequency from their native frequency due to blending of polymer resulting in the formation of weak hydrogen bond.

**Figure 2.**
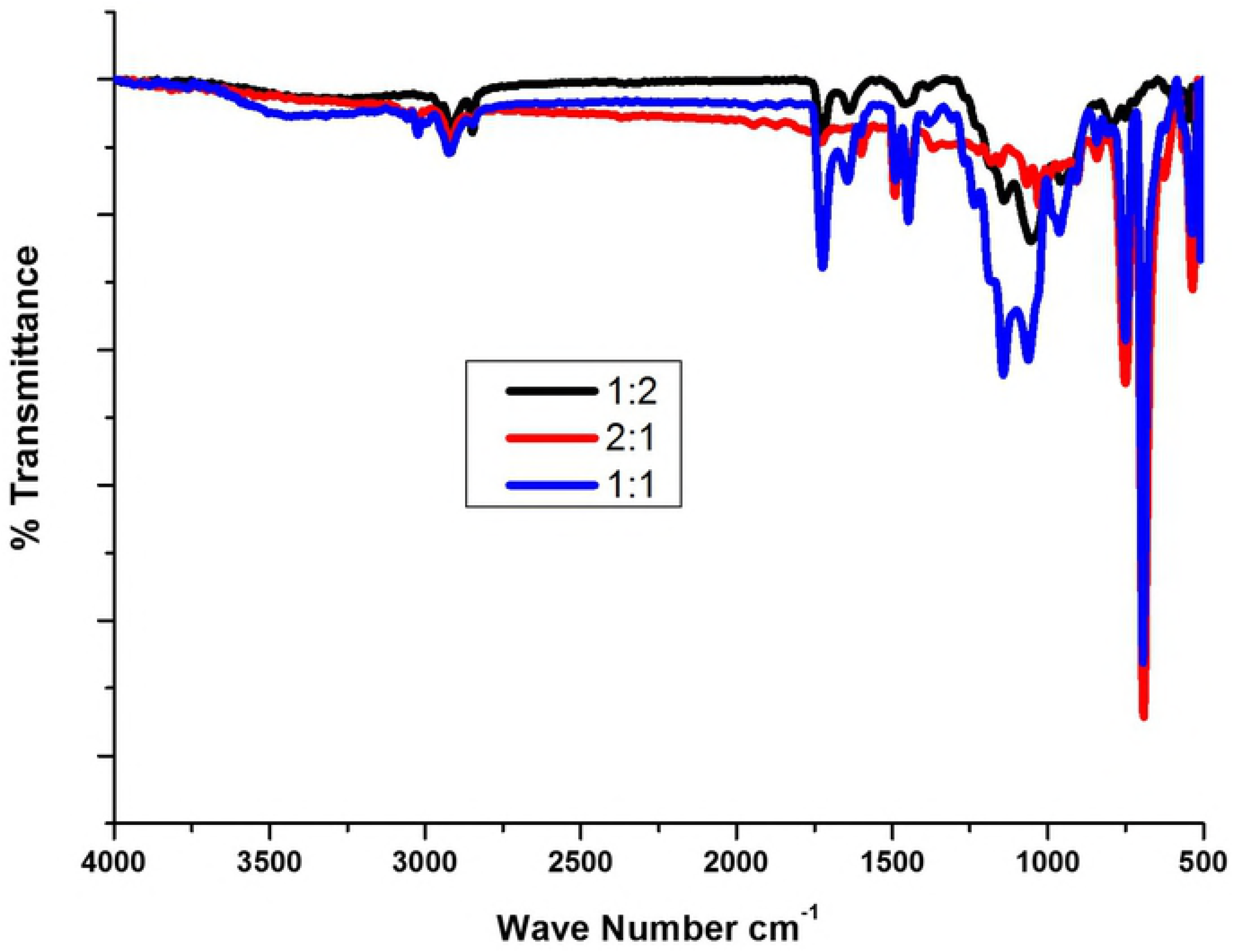
FTIR analysis of showing the interaction between PMMA and PS.

The blend containing 1:1 ratio showed characteristic absorption peaks of PMMA corresponding to asymmetric stretching of CH_3_ and C=O were assigned to bands 2951 cm^-1^ and 1736 cm^-1^ (Figure 2). The peaks 1482 cm^-1^ is a characteristic vibrational band for CH_2_ scissoring, similarly the vibrational band at 1452 cm^-1^ corresponding to asymmetric vibration of CH3 stretching or due to the PMMA deformation which was further confirmed by OCH3 deformation peaks at 1390 cm^-1^ (Figure 2). In addition, the absorption peaks at 1600 and 698 cm^-1^ in all the blends represents the C-C stretching and ring deformation of polystyrene. The broadened peaks at 750, 1050 and 1200 were observed for blend of 1:1 which are correspond to CH_2_ twisting, wagging and rocking respectively, whereas the blend of 2:1 and 1:2 appeared to be smooth curve in the same region [24]. The IR results confirm that the blend was strongly mixed and formation of polymer blends in case of equal concentration of polymer compared to other two concentrations. It was observed that the results were in par with other reports and confirmed that there was no major shift in the peaks hence the blends may be due to physical bonding rather than a chemical one [25].

### Water absorption studies

**Figure 3:**
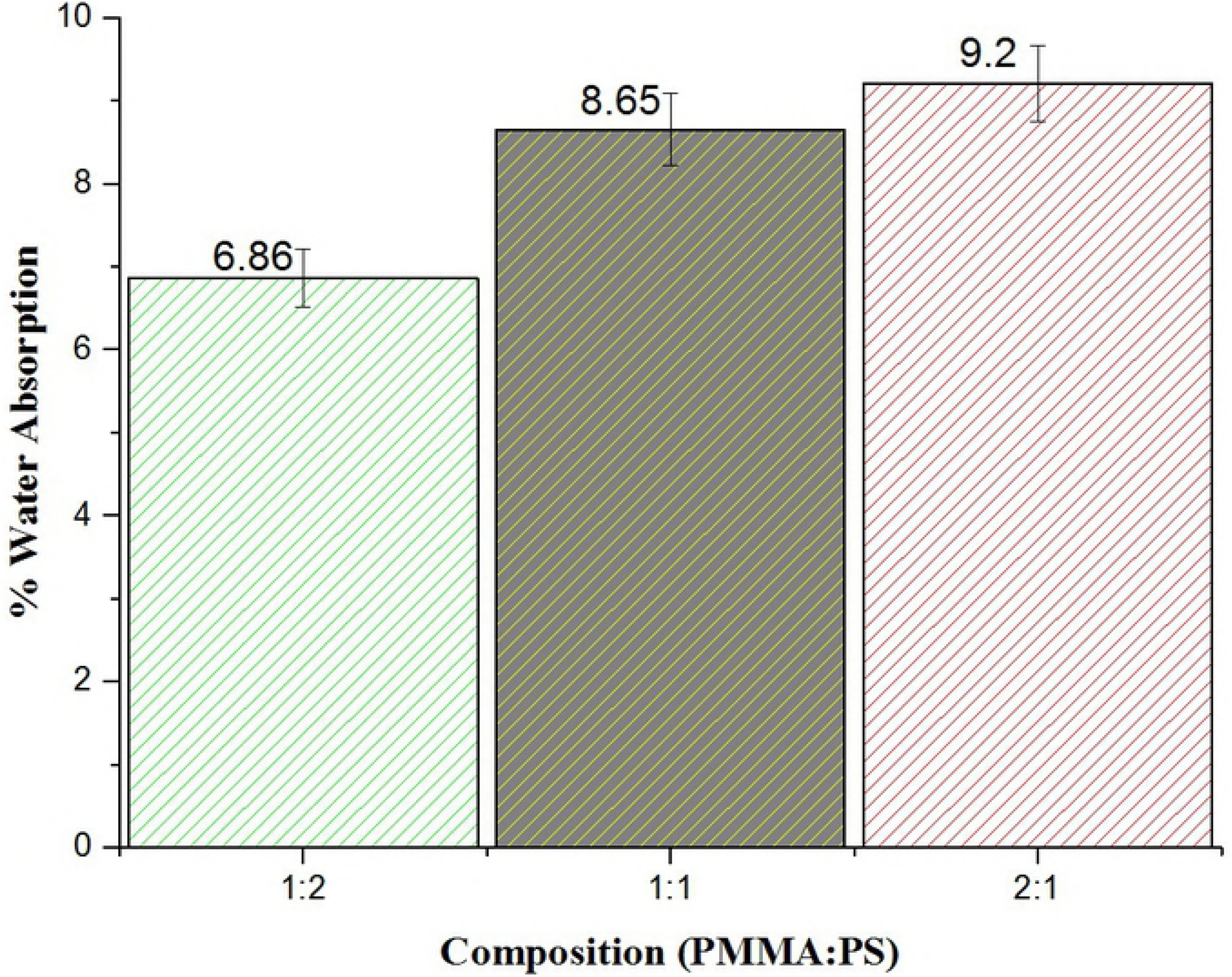
Percentage water absorption of the prepared composite membranes.

The results of the water absorption studies for the three groups is shown in Figure 3. Figure 3 shows that the composite membrane of PMMA/PS 2:1 has the maximum water absorption while the membrane of 1:2 has the least water absorption. This variation is attributed to the hydrophobicity of polystyrene. Hence, it was observed with increase in concentration of polystyrene decrease in water absorption capacity of the prepared composite membranes. The swelling capacity of membranes increases with decrease in the dimensional stability, which in turn directly influence the release of drug from membrane.

### Surface Morphology – SEM Analysis

The SEM images of the composite samples are shown in Figure 4. From the images, it was clearly seen that in all the samples, the silica fillers are evenly distributed throughout the polymer matrix.

**Figure 4.**
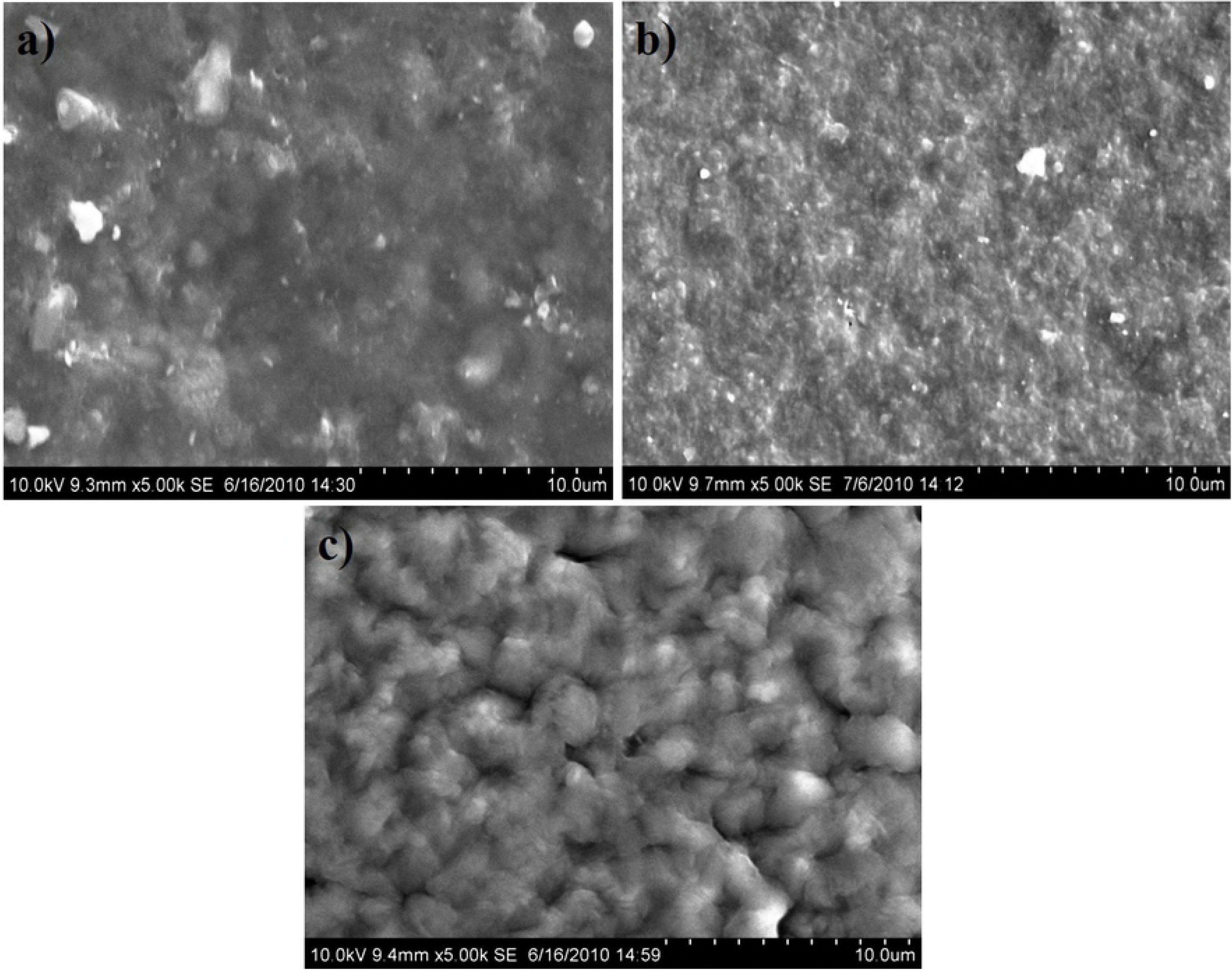
SEM images of a) PMMA:PS (1:2) b) PMMA:PS (1:1) c) PMMA:PS (2:1)

The cracks that appear on the surface of the composite membranes are due to capillary pressure occurring during the solvent evaporation. The presence of these cracks on the surface will be advantageous for drug delivery and tissue engineering applications. The presence of these pores and cracks on the surface will help in the ingrowth of the hydroxyapatite crystals thus enabling good mechanical and chemical bonding of the bone with the polymer surface.

### Tensile strength

The tensile strength of the prepared samples is shown in figure 5. From the figure, it is seen that the sample of sub group 3 having PMMA: PS ratio of 2:1 has the maximum tensile strength of 78.3 MPa. With a decrease in the concentration of PMMA in blend membrane, the tensile strength is also found to decrease gradually due to elastic nature PMMA possess.

**Figure 5.**
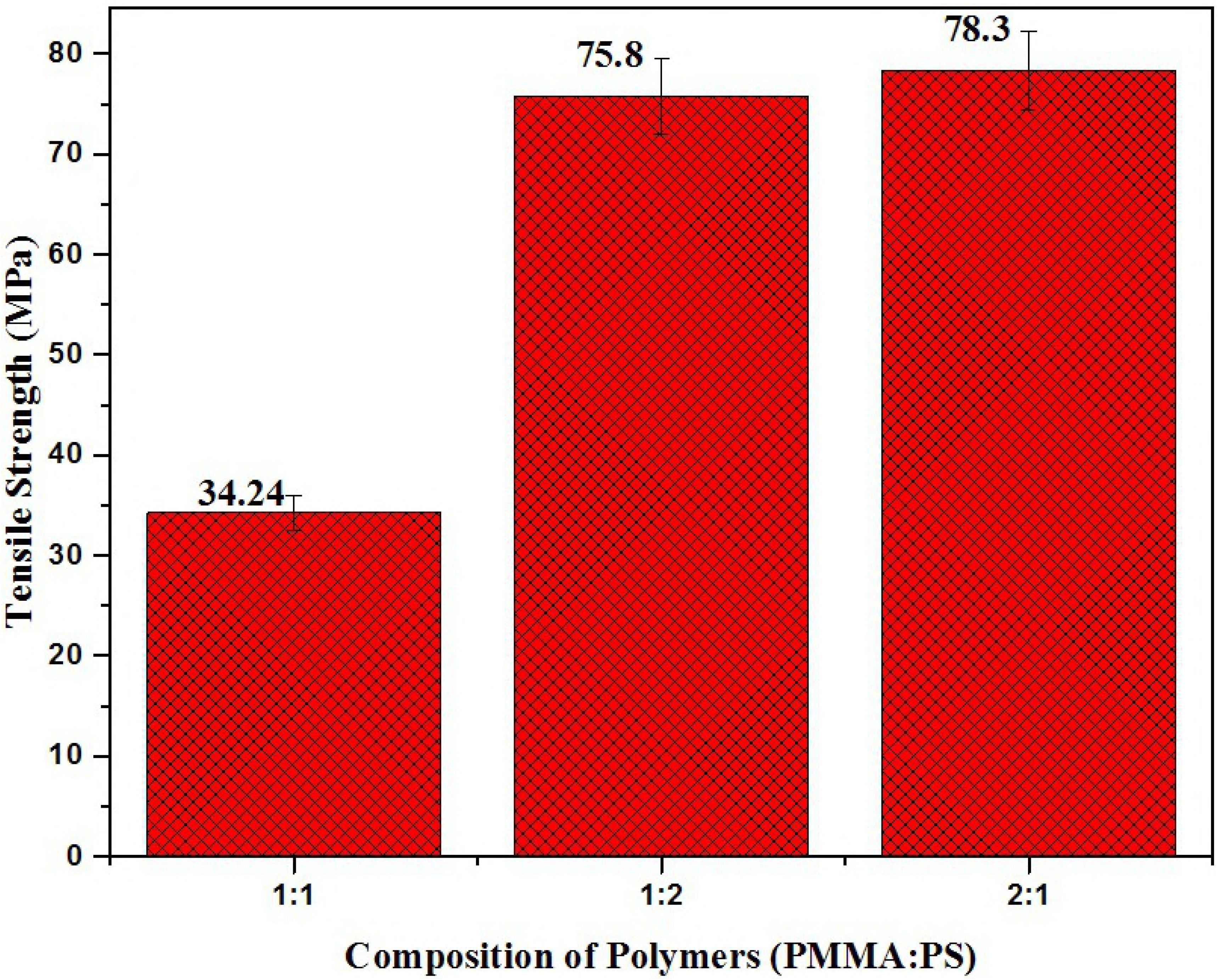
Tensile strength of the prepared composites.

### Bioactivity study

Bioactivity of the prepared composite membranes was determined by keeping the membranes immersed in Kokubo solution and observing the growth of the apatite under the scanning electron microscope. The SEM images are shown in figure 6

**Figure 6.**
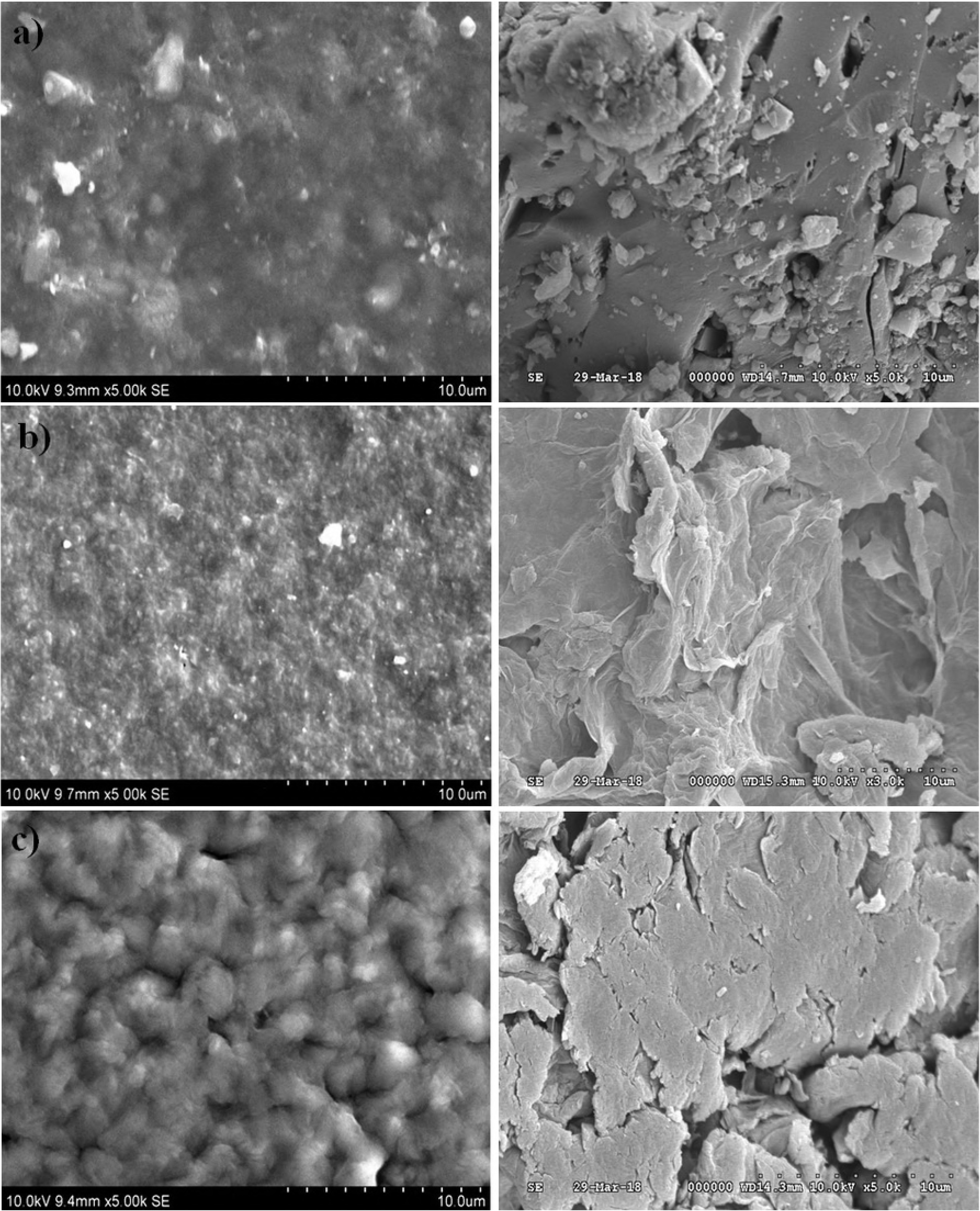
Shows the apatite growth on the surface of the subgroups 1, 2, 3 of PMMA:PS with silica fillers.

The SEM images of the three subgroups after immersion in SBF for 15 days at room temperature is shown in figures 6 that correspond to with and without the silica fillers in composites. It was clear from figure 6 a, b, c that all the composites incorporated with si fillers showed excellent bioactivity compared with to the ones that were not incorporated with si. It was clear from the SEM that the si enhance the bioactivity of membranes and it was observed that PS as well plays an important role in the bioactivity. It was observed from the figure 6a, b & c that the composites with equal concentration of PMMA and PS were highly bioactive and the composite with more wither of PMMA or PS showed comparatively lower amount of bioactivity. The bioactivity of composites with more concentration of PS was less bioactive compared to other two composites respectively.

### Drug Release Kinetics

The composites prepared for applying as bone cement was checked for its effective drug release kinetics as it may serve for multipurpose in enhancing the bone growth with active release of drug (Figure 7). It was observed from drug release kinetics that the drug was released in a sustained manner for a period of 180 h in PBS at pH 6.8. The initial phase or burst release was observed in all three cases the models that is in par with other research reports [26–29]. It has to be noted that the drug release in model 1 may be due to fickian diffusion and took a very long time owing to higher concentration & hydrophilic nature of PMMA that makes the drug release in a much-sustained fashion. This was not the case with model 2 & 3, which exhibited a quick release of drug compared to model 1, and this can be attributed to the complete degradation between the blended membranes since it was not a chemical bonding between the two polymers. The release kinetics were in par with FTIR results and confirmed the presence of physical interaction between polymers that might have resulted in quick release of drug from composites.

**Figure 7.**
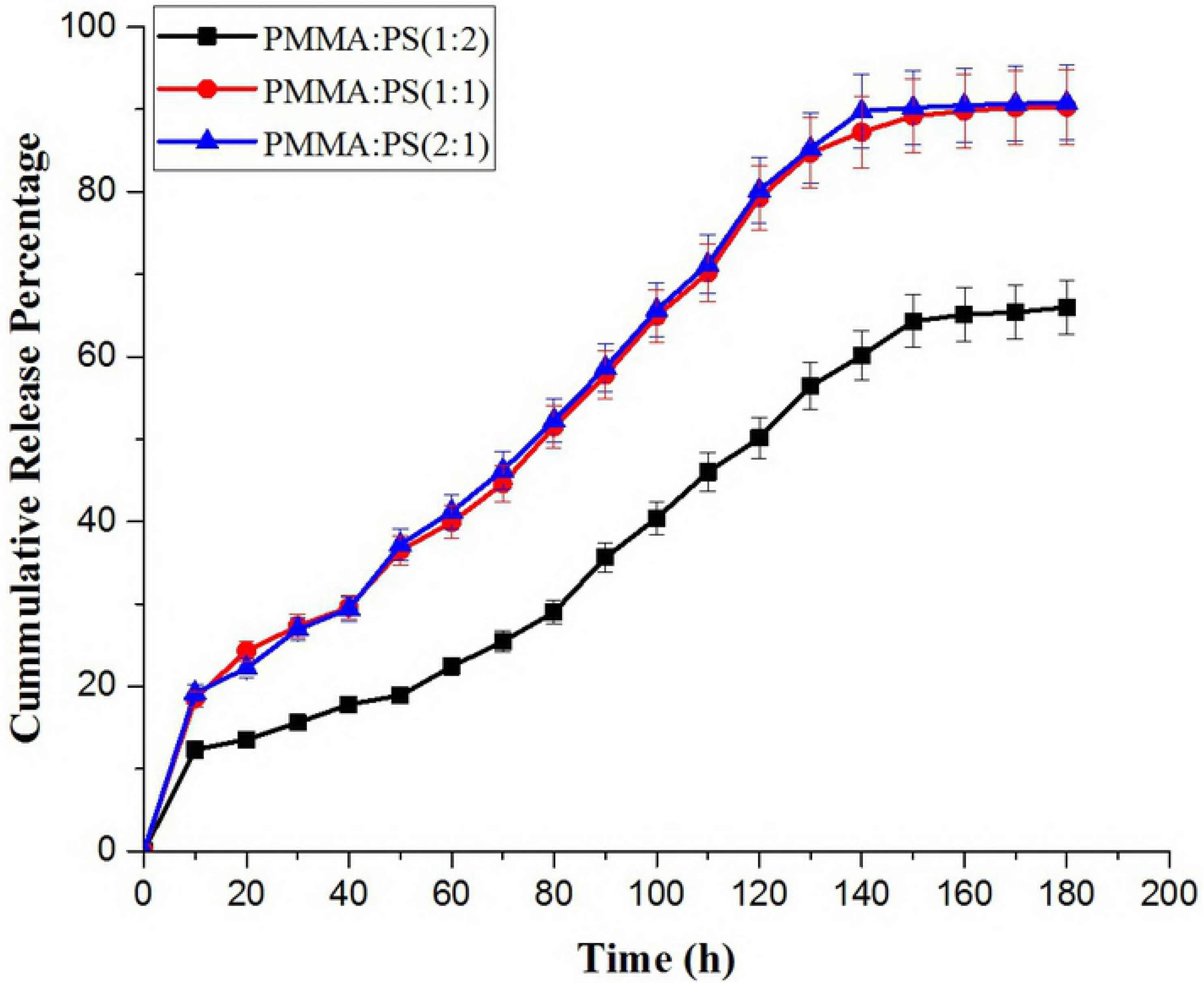
shows the drug release kinetics of the prepared composite materials with varying concentration of PMMA: PS.

The release kinetics were fitted onto peppas model to further be confirmative about the type of release that might have occurred from composite models [30]. The model 1 had a value of n≥5 when fitted onto peppas model attributing to the fickian diffusion, the other two composite did not fit into any other models, which is out of the scope of our study.

### *In vitro* Cytotoxicity by MTT method

The cytotoxicity studies performed showed interesting results. Figure 8 shows the RAW macrophage 264.7 cells after treating with composite membranes for 48 h. It can be observed that the cells treated with equal concentration of PMMA: PS showed confluent cells compared with the other two concentrations. It can be visually observed that amount of viable cells very lower in case of higher concentration of either PMMA or PS. Figure 8 (b) shows high density of macrophage cells indicating its compatibility for cell proliferation.

**Figure 8.**
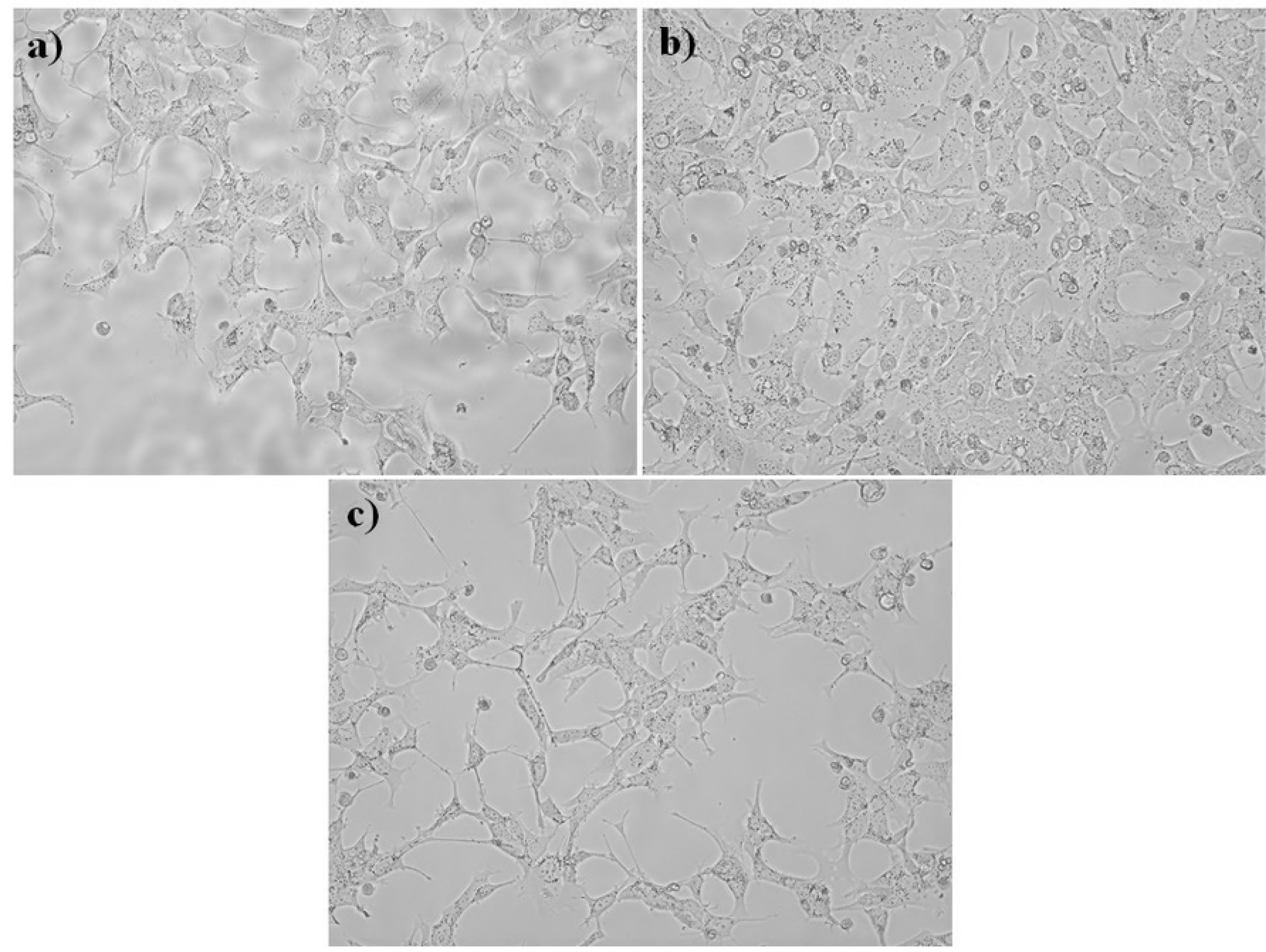
Microscopic images of cytotoxicity study. a) Model 1, b) Model 2, c) Model 3.

The results were in par when studied with MTT assay where the composites with equal concentration of polymers showed an enhanced viability compared to the other two models. The percentage viability of the cells was calculated after noting the OD values of the control. The results are displayed in figure 9 in which a graph is plotted by taking the cell count of the control group as 100 percentage. The viability percentage was in the range of 93.5±3 in case of equal concentration of polymer whereas it was lower on the other two cases. Among the three groups, the sub group 2 has the best biocompatibility among other subgroups of PMMA: PS.

**Figure 9.**
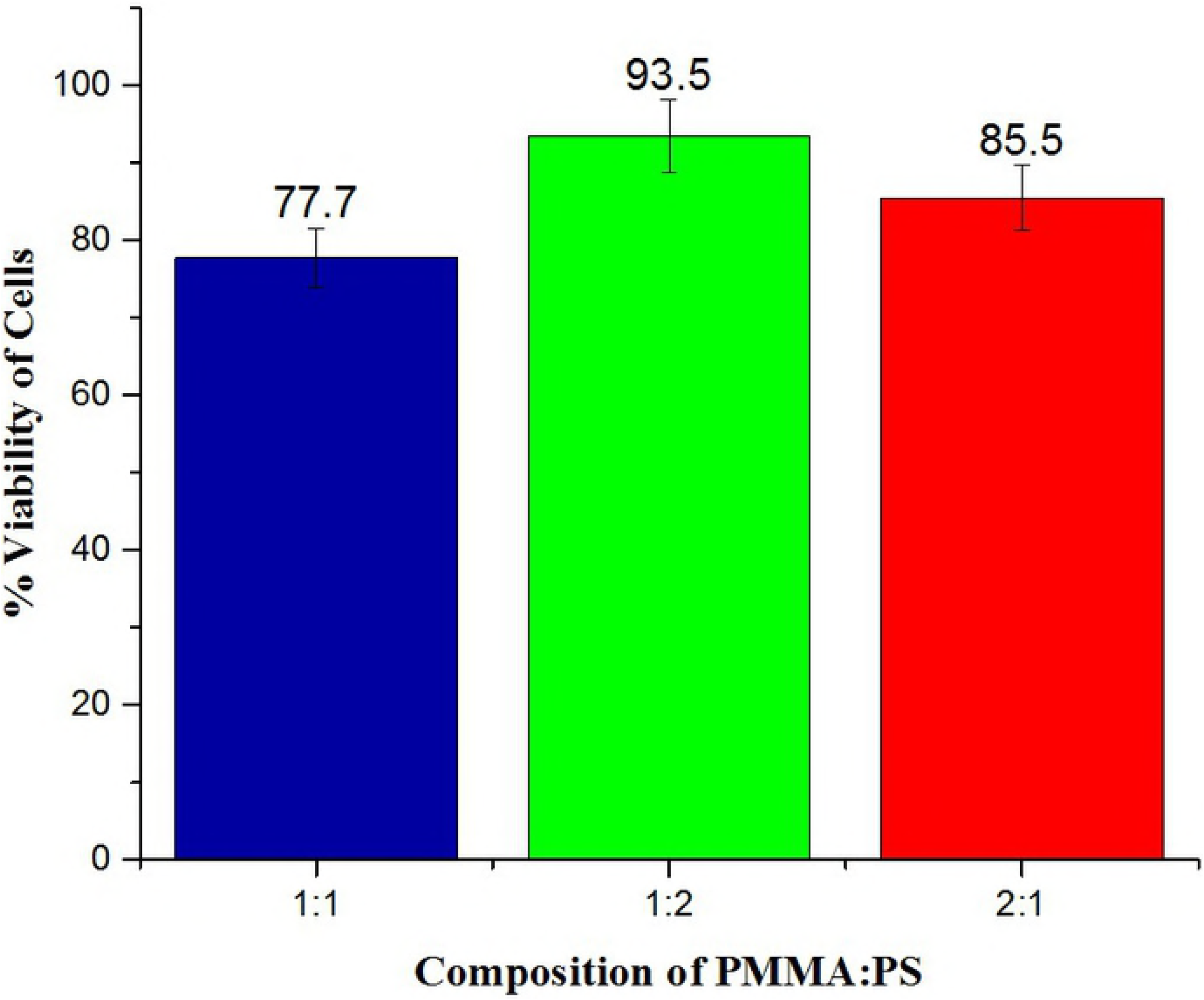
Graphical representation of the cell viability of the three groups.

### *In vitro* Inflammatory Response

The importance of these composites when applied *in vivo* has to be taken into consideration since most of them cause inflammation when applied in vivo or during the application as prosthesis. Hence, in the present study, we evaluated inflammatory response of these composites on RAW macrophage cells and its effect was analyzed with inflammatory specific genes like TNF-α, IL-6 and IL-1 (figure 10). It was observed from the results that most of composites were not causing a major inflammation in all the three cases and in particular the expression was least in case of equal concentration of PMMA and PS. It was also observed that the expression of IL-6 marker gene was at a higher end in case of higher concentration of PMMA. The results from qPCR confirmed that the concentration has a direct role in causing inflammation particularly in dental aplications that is in par with the one reported by Spasojevic et al [14] as they. It can be observed that these onflammatory expression was at lower end with the addition of PS only at a particular concentrations. When the concentration of PS was increased the inflammatory resposne as well got increased slightly compared to the equal concentrations. Hence, it is evident that only at equal concentration of PMMA and PS the composites very good properties. The increase in concentration of PMMA or PS was not good property as that of the equal concentration. Therefore, the composites with equal concentration of PMMA-PS can eb suitable for bio medical apllications.

**Figure 10.**
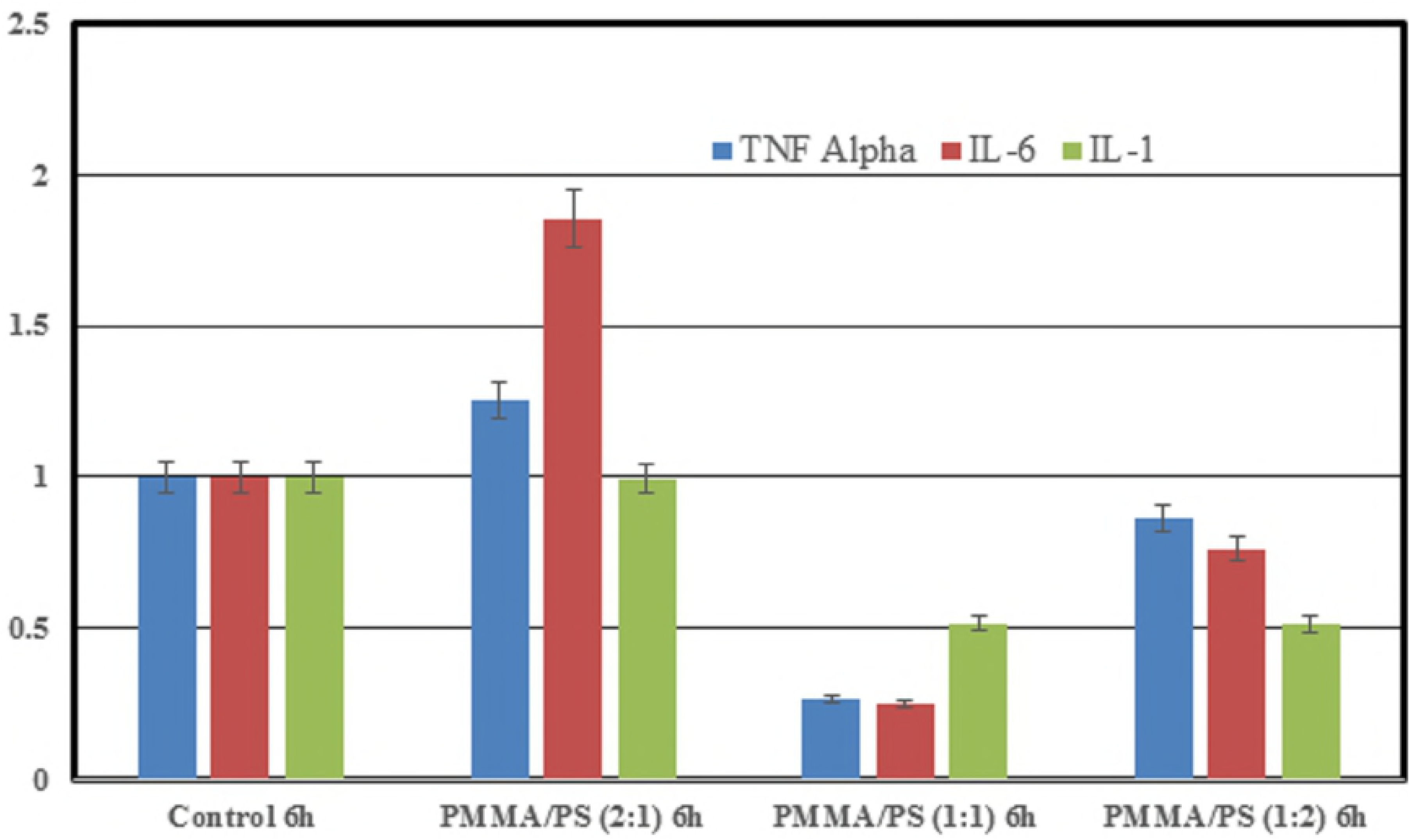
Inflammatory response observed on RAW macrophage cells – *in vitro*.

## Conclusion

Polymer silica composites of PMMA/PS were successfully fabricated by sol-gel technique. SEM and FTIR characterized the blend membranes and confirmed the presence of physical bonding between them. The SEM images showed that the surface homogeneity of the membranes was dependent on the concentration and blending between the two constituent polymers. The blend formation between the polymers in the three groups was confirmed by FTIR. The bioactivity of membranes were analyzed by immersing the samples in SBF and examining the surface for the formation of apatite layer under the scanning electron microscope (SEM). Among the three groups, model 1 containing equal concentration of PMMA/PS blend composite showed the high bioactivity while the group two showed the most dense apatite formation of Ca-P crystals on its surface. The cytotoxicity studies were performed to evaluate the biocompatibility of the fabricated composite membranes. The *in vitro* study carried out by MTT method showed the composite with equal concentration of group II and III samples exhibited excellent cell viability while the group I, the sample of the subgroup 3 namely PMMA:PS in the ratio of 2:1 exhibited favorable cytotoxicity value than the other two subgroups. From this study, it can be concluded that the sol-gel technique is versatile method to fabricate the polymer composites having immense biomedical application. Among the polymers investigated, model 1 samples were less suited for load bearing orthopedic applications but more suited for sustained drug delivery and tissue engineering application. Model 1 showed very good tensile properties whereas other two models showed optimum biocompatibility and moderate strength that shall be suitable orthopedic application.

## Acknowledgement

The first author acknowledges the support for performing cytotoxicity assay from Professor Szondy Zsuzsanna, Apoptosis Laboratory, Department of Biochemistry & Molecular Biology, University of Debrecen, Hungary and UGC Stipendium Hungaricum (Educational Exchange) Programme. The first author would also like to thank DST for providing fellowship and support vide Lr. No. DST/INT/Korea/P-47 Dated 18.05.18. The authors would like to thank Professor Csaba Cserháti, Department of Physics, University of Debrecen for his kind support in carrying out Scanning Electron Microscopy Analysis.

